# First outbreak of Lumpy Skin disease in Catalonia, Spain, 2025-2026

**DOI:** 10.64898/2026.06.18.733166

**Authors:** Pau Obregon-Gutierrez, Florencia Correa-Fiz, Osvaldo Fonseca-Rodríguez, Martí Cortey, Àlex Cobos, Carles Riera, Mercè Soler, Núria Ribas, Francisco Domenes, Lola Pailler-García, Mariano Domingo, Natàlia Majó, Enric Vidal, Cristina Lorca-Oró

## Abstract

Lumpy skin disease (LSD) is an emerging cattle disease caused by lumpy skin disease virus (LSDV), with major impacts on the industry, being classified as a Category A disease. Although it was historically confined to Africa, LSD has expanded into the Middle East, Asia and Europe. Here, we report two LSDV genomes from the first outbreak detected in Catalonia, Spain, in October 2025. The genomes were assembled from high-throughput sequencing data generated from two homogenized skin nodules. Comparative phylogenetic analyses were performed using all available complete LSDV genomes and *rpo30* gene sequences. These analyses placed the LSDV isolates detected in Catalonia within clade 1.2, closely related to the isolates recently reported in Sardinia, Italy. Our findings also support a connection between recent south-western Europe and central African strains, possibly through northern Africa, and highlight the need for more complete genomes to clarify the origin and connections among recent LSDV outbreaks.

## Introduction

Lumpy skin disease (LSD) is an emerging transboundary disease of cattle and water buffalo caused by *Capripoxvirus lumpyskinpox* also known as lumpy skin disease virus (LSDV). The disease is characterized by nodular skin lesions, fever, respiratory signs, edema, lymphadenitis, reduced milk production, and infertility (1). Although other ruminant species can be infected by LSDV, their epidemiological role and clinical relevance remain incompletely understood. Because of its major socioeconomic impact associated with decreased productivity, animal welfare, and trade restrictions, LSD is classified as a Category A disease under the European Animal Health Law and is a notifiable disease to the World Organization for Animal Health (WOAH). Consequently, immediate eradication measures are required, including stamping out of affected herds, strict movement restrictions and emergency vaccination.

LSDV transmission occurs seasonally mainly through mechanical spread by hematophagous arthropod vectors, especially the stable biting fly (*Stomoxys calcitrans*) but also including mosquitoes, Culicoides, tabanidae and ticks, which play a critical role in virus maintenance and dissemination in short distances (1). Human driven movement of live animals largely contributes to long distance spread of the disease (2,3).

Historically confined to Africa, LSD has expanded during the last decade into the Middle East, Asia and Europe, reaching south-eastern Europe in 2015 (4). Large-scale vaccination campaigns and depopulation of affected herds have contributed substantially to disease control in affected regions; however, the recent detection of LSDV in Italy, France and Spain in 2025 indicates that new incursions into Europe continue to occur a decade after the first south-eastern European epidemic (3,15). Here, we report the first complete genomic sequences of two LSDV isolates detected in Catalonia, Spain, in October 2025, and investigate its phylogenetic relationships, genetic variability, and possible epidemiological origin through comparison with publicly available global genomes. This study provides novel molecular data that contribute to understanding the first documented introduction and spread of LSDV in Spain.

## Methods

### LSDV sampling and processing

Two skin nodules collected from naturally LSDV-infected cattle from two independent affected farms in Catalonia (Spain) sampled the 7^th^ and 9^th^ of September 2025 were individually homogenized in sterile phosphate-buffered saline (PBS) using mechanical disruption. Total DNA was extracted from the resulting tissue homogenates following the manufacturer’s instructions (IndiMag Pathogen Kit, Indical Bioscience). The presence of LSDV was confirmed by specific PCR (Bio-T kit® Lumpy Skin Disease, Biosellal). To increase the proportion of viral DNA, host DNA depletion was performed using the NEBNext® Microbiome DNA Enrichment Kit (New England Biolabs, USA) according to the manufacturer’s protocol. The two samples were identified as LSDV_CAT2025_5380 and LSDV_CAT2025_9248, respectively.

### DNA sequencing

Prior to library preparation, DNA samples were purified using magnetic beads (NEB E2612 protocol). DNA quality was assessed by spectrophotometric evaluation of the 260/280 and 260/230 absorbance ratios, and DNA concentration was determined by fluorimetry using the Qubit system (Thermo Fisher Scientific, USA).

Sequencing libraries were prepared using the Illumina DNA Prep kit (Illumina, USA) and library quality was evaluated using the Agilent TapeStation High Sensitivity system (Agilent Technologies, USA). Paired-end sequencing (2 × 150 bp) was performed on an Illumina NextSeq 1000 platform. Raw sequencing data were demultiplexed to generate FASTQ files.

### Bioinformatic analysis

In order to retain only reads belonging to LSDV, reads were initially filtered using kraken2 v2.1.3 (5) against its viral database (available in: https://benlangmead.github.io/aws-indexes/k2;accessed on May 2026). After that, all reads classified as *Capripoxvirus lumpyskinpox* (taxid 3427718) were retained using kraken tools (https://github.com/jenniferlu717/KrakenTools). In addition, reads were filtered using fastp v1.1.0 (6), with automatic adapter detection, a quality threshold of 20, polyG/polyX trimming and a minimum read length of 75 bp after filtering.

*De novo* assembly was performed using a subsampled set of 100,000 paired-end reads to reduce both excessive sequencing depth and duplicated read representation, while maintaining sufficient coverage for viral genome reconstruction. Reads were subsampled using seqtk v1.4-r122 (https://github.com/lh3/seqtk) with a fixed random seed, and the resulting read set was assembled with SPAdes v4.0.0 (7) using the --careful option and multiple k-mer sizes. Scaffolds obtained from the *de novo* assemblies were aligned to the LSDV reference genome (NCBI acc. AF325528.1) using minimap2 v2.28-r1209 (8). Since the repetitive terminal regions of the viral genome could not be unambiguously resolved by *de novo* assembly, a map-based genome assembly complementary approach was subsequently performed.

For the map-based assembly, reads were mapped against the reference genome using BWA v0.7.18-r1243 (9). Alignments were sorted and indexed, and mapping statistics and genome-wide coverage were obtained with SAMtools 1.21 (10). Positions with coverage below 10X were masked as Ns to avoid unsupported consensus calls. Variants were called with bcftools v1.21 (11) under a haploid model (--ploidy 1), aiming to recover the dominant viral consensus sequence present in each sample. Consensus genomes were generated with bcftools (11), aligned to the reference using MAFFT v7.525 (12) and compared with snp-sites v2.5.1 (13). Average nucleotide identity (ANI) values between genomes were calculated using pyani v0.3.0 (14).

Two different approaches were followed to perform the phylogenetic analyses. The main phylogenetic analysis included all available complete LSDV genomes (WGS-phylogeny) downloaded from NCBI (https://www.ncbi.nlm.nih.gov/datasets/genome/?taxon=59509, accessed on May 26^th^ 2026). All partial/incomplete genomes and/or with < 150,000bp length were excluded from the analysis. Three recently published Italian genomes (15) were manually added to the collection (since they were not included in NCBI-genome database), up to a final pool of 122 genomes. These genomes were aligned using MAFFT v7.525 (12), and the multiple alignment was trimmed with trimal v1.5 (16) using --automated1, a heuristic selection of the automatic method based on similarity statistics (optimized for Maximum Likelihood phylogenetic tree reconstruction). The maximum likelihood phylogenetic tree was built using IQtree v3.1.0 (17) with 1,000 replicates for ultrafast bootstrap (-B) and extended model selection followed by tree inference (-m MFP). Another phylogenetic analysis was done using only the sequences included in Fadele, *et al* (18), including the two strains from Cameroon reported in the study, which was performed using the same procedure described above.

In addition, we also performed a gene-based phylogenetic analysis using *rpo*30 gene. Here, all sequences belonging to *rpo*30 gene from LSDV were downloaded from nucleotide NCBI database (accessed on May 28^th^, 2026). Additionally, we annotated the complete genome sequences from the WGS-phylogenetic analysis described above using Prokka 1.15.6 (19) to extract the *rpo*30 genes. The combined collection including 486 *rpo*30 sequences was then aligned, trimmed and used for maximum likelihood phylogenetic reconstruction following the same pipeline described for the WGS-analysis.

Variant calling was performed using the genomes belonging to the clade under study. Variants were identified with Snippy v4.6.0 (https://github.com/tseemann/snippy), using the assembled genomes as contig inputs. The predicted functional effects of the identified SNPs were annotated with SnpEff v5.0 (20).

All FASTQ files were quality-checked using FastQC v0.12.1 (21), and basic read statistics were monitored with SeqKit v2.10.1 (22). Genome assemblies were evaluated using QUAST (23), while FASTA files were inspected and processed with SeqKit v2.10.1 (22). Sequence alignments were visually inspected using AliView v1.30 (24). Phylogenetic trees were visualized and processed in iTOL v7 (25). Output files generated by the different tools were processed with in-house Bash and R scripts. R scripts were run in R v4.5.2 (26) within RStudio version 2026.5.0 (27), using the tidyverse (28) and ggplot2 (29) packages for data processing and figure preparation.

The full pipeline can be found in: https://doi.org/10.34810/DATA3412. The genome sequences are publicly accessible in Genbank under accessions PZ500253 and PZ500254 for strains LSDV_CAT2025_5380 and LSDV_CAT2025_9248, respectively. The raw reads can be found in NCBI BioProject PRJNA1475247, SRA accessions SAMN60634786 and SAMN60634787.

## Results

Samples from the two homogenized skin nodule samples were processed and tested positive for specific LSDV PCR, with a Ct of 20 and 19 for isolates LSDV_CAT2025_5380 and LSDV_CAT2025_9248, respectively. After host DNA depletion, total DNA was submitted to NGS sequencing. Raw reads were processed and filtered against the kraken2 viral database in order to retain the reads of the responsible LSDV strain. After quality and length cleaning, the final number of reads was 531,174 and 3,700,045 for isolates LSDV_CAT2025_5380 and LSDV_CAT2025_9248, respectively (**Table 1**).

**Table 1.** Number and percentage of reads: from initial reads to classified as non-viral or *Capripoxvirus lumpyskinpox*, and the final reads after filtering by quality and length.

| Reads | LSDV_CAT2025_5380 | LSDV_CAT2025_9248 |
| --- | --- | --- |
| Initial | 30,419,156 | 23,120,766 |
| Non-viral | 29,594,076 (97.29%) | 18,363,900 (79.43%) |
| *Capripoxvirus lumpyskinpox* | 589,612 (1.94%) | 3,859,052 (16.69%) |
| Filtered reads | 531,174 (1.75%) | 3,700,045 (16%) |

*De novo* assembly of the final filtered reads yielded two scaffolds for both samples, with identical sizes: a main scaffold of 145,951 bp and a smaller scaffold of 2,438 bp, for a total assembled length of 148,389 bp. Alignment against the LSDV reference genome (AF325528.1) showed that the main scaffold covered most of the genome, whereas the smaller scaffold aligned to both terminal regions, consistent with unresolved repetitive ends. No additional scaffolds suggesting sequences absent from the reference genome were identified. Therefore, the complete viral genome could not be fully reconstructed by *de novo* assembly alone, and a reference-based assembly strategy was selected for final genome reconstruction and phylogenetic inference.

Reference-based assembly generated a single-scaffold consensus genome for each sample. The final sizes were 150,787 bp for sample LSDV_CAT2025_5380 and 150,768 bp for sample LSDV_CAT2025_9248, similar to the lengths of the available LSDV genomes. Read mapping rates were very high for both samples, with 99.96% and 99.97% of reads mapped and a mean coverage of 985.3X and 7,119.4X respectively, with no positions showing zero coverage. Nevertheless, after masking positions with low coverage (<10×), the final consensus sequences contained 93 Ns for LSDV_CAT2025_5380 and 1 N for LSDV_CAT2025_9248, most of them in both ends of the genome. These reference-based consensus genomes were used for downstream comparative analyses.

WGS-phylogeny using all available complete LSDV genomes (**supplementary Table 1A**) recovered the major genomic structure previously described for LSDV, clearly separating subgroup 1.1, corresponding to the Neethling-like clade and Asian variants, from subgroup 1.2, which includes Kenya-like and recent wild-type viruses, as previously reported (30,31). The two genomes analysed in this study, were placed within LSDV clade 1.2 (**Figure 1**). More specifically, they grouped within the same phylogenetic region as the Italian genomes reported in 2025 and showed close proximity to the Nigerian LSDV isolate V281 from 2018 (NCBI accession OK318001.1), in agreement with the recent Italian study (15). Besides, the phylogeny did not provide sufficient bootstrap support to define the two genomes as distinct phylogenetic sublineages. Consistently, the ANI between both isolates was 99.98%, indicating a very high genomic similarity. In addition, comparisons with the Italian genomes also showed high ANI values, around 99.8% or higher (range 99.79-99.98%), supporting the close relationship with these strains. The same phylogenetic placement was obtained when the analysis was repeated using the main contig of 145,951 bp generated by *de novo* assembly.

**Figure 1.**
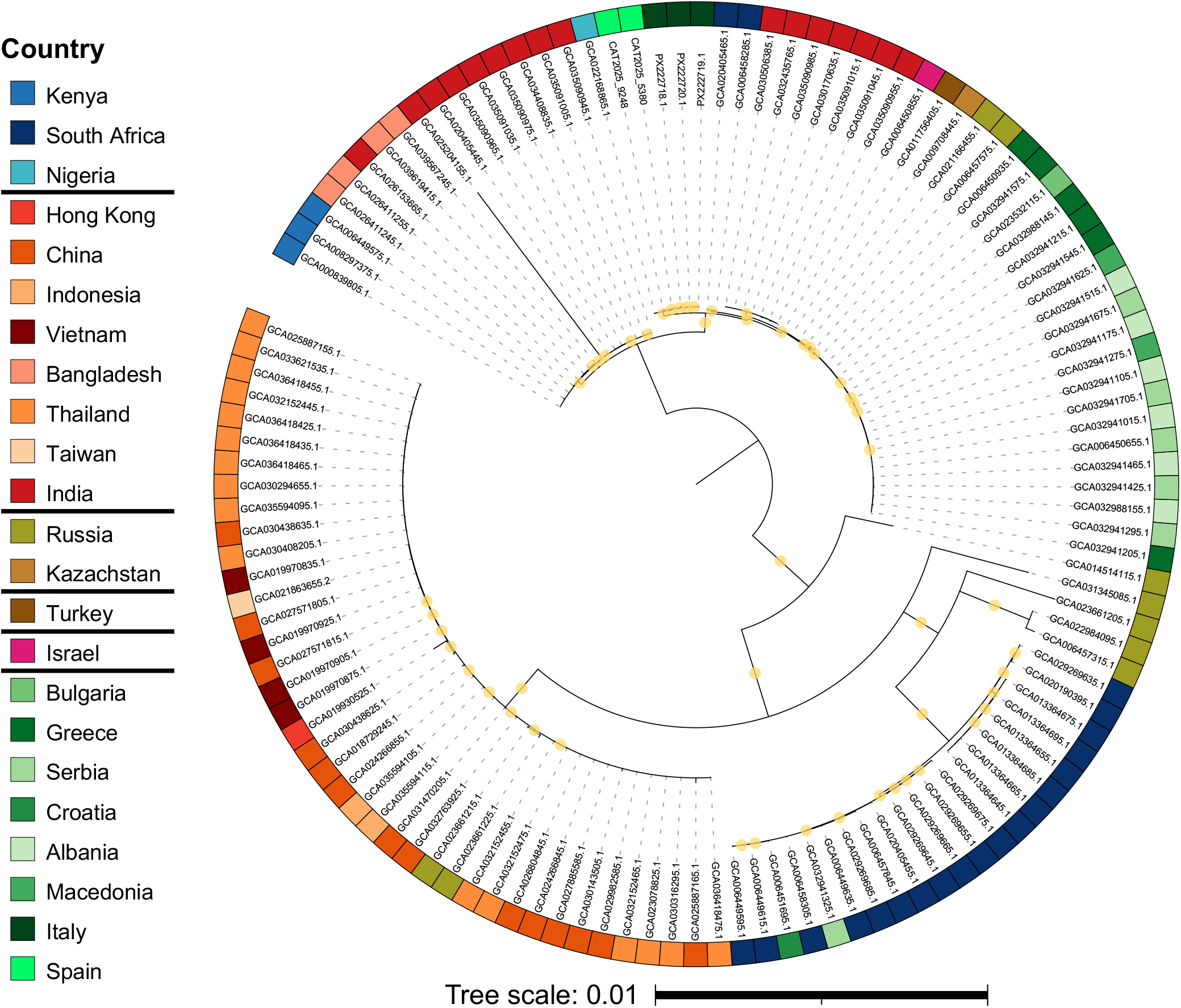
Maximum likelihood phylogenetic tree of all available complete LSDV genomes. The colour code corresponds to the site of isolation, with similar colours within each continent or geographical area. The Goatpox virus Pellor (NCBI acc. NC_004003) was used as outgroup but was removed to improve visualization of the clusters and the tree was re-rooted at midpoint. Yellow circles indicate branch bootstrap support equal or greater than 70. Branches with bootstrap support□<□70 have been collapsed. The names of the strains characterized in this study are shown, while the NCBI accession is provided for all others.

In addition, we performed a different phylogenetic tree to further corroborate the phylogenetic relationship and placement of the Spanish strains. Since Fadele, *et al*. recently characterized two Cameroonian LSDV strains, and observed that these strains also clustered with the Nigerian isolate V281 suggesting that these may represent a distinct sub-lineage within LSDV clade 1.2 (18), we reproduced the phylogeny with the same subset of genomes including the new Spanish and Italian genomes. This phylogenetic reconstruction reproduced the clade 1.2 topology reported in that study, with the Cameroonian genomes grouped with the Nigerian isolate V281, supporting their proposed placement within a distinct sub-lineage of LSDV clade 1.2. Importantly, the Italian and the Spanish genomes were also placed within this same phylogenetic group (**Figure 2**).

**Figure 2.**
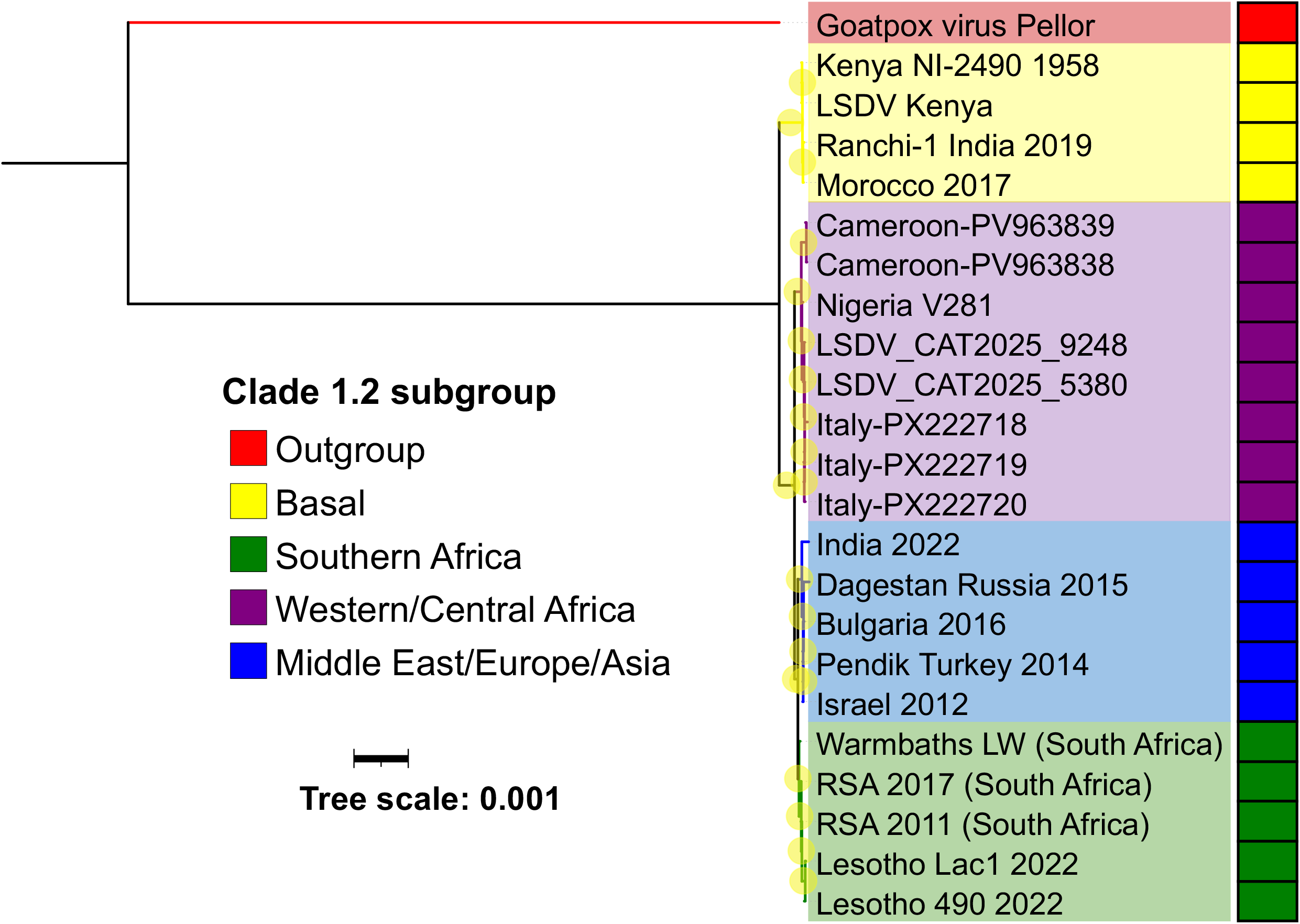
Maximum likelihood phylogenetic tree of selected genomes. Reconstruction of the phylogenetic tree reported by Fadele, *et al*. (2025) including the Nigerian V281 sequence and partial sequences form Cameroon, together with Spanish and Italian strains. The tree was rooted at the outgroup (Goatpox virus Pellor NC_004003, indicated in red). Yellow circles indicate branch bootstrap support equal or greater than 70. Branches with bootstrap support□<□70 have been collapsed. The colour code corresponds to LSDV clade 1.2 subgroups described in the stated study.

To broaden the representation of LSDV diversity, particularly from African lineages, an additional phylogenetic analysis was performed using the *rpo30* gene, including 486 sequences (**supplementary Table 1B**). This gene has been widely used in previous LSDV phylogenetic studies and is considered informative for assessing genetic relationships among LSDV strains (32–37). This allowed the inclusion of 107 additional African strains, including more Nigerian sequences. Although the low variability of this marker limited the resolution of many internal branches, a supported cluster including the Spanish, Italian together with some Nigerian sequences was recovered (**supplementary Figure 1**), consistent with the separation of the Spanish/Italian lineage from the southeastern European LSDV strains. Regarding the two Cameroonian sequences, one clustered with the majority of African strains within a distinct subclade, whereas the other appeared more isolated, possibly reflecting some ambiguity in the *rpo30* region.

In order to characterize the differences accumulated in the recent south-western European LSDV strains, the two Spanish and three Italian genomes were compared with the closely related African strain (Nigerian V281), used as the genomic reference. This reference-based comparison allowed the identification of SNPs, insertions and deletions distinguishing the new European genomes from the Nigerian strain (**Figure 3**). Overall, the Spanish and Italian genomes showed a highly similar pattern, with many variants shared across the five European genomes relative to the African genome. Many variants were either synonymous or occurred in non-coding regions. However, several non-synonymous changes were also detected, including missense mutations and short length indels (1-3 nucleotides) affecting coding sequences. Some of the non-synonymous variants affected genes annotated as interferon gamma receptors, kelch-like proteins, ankyrin-like proteins, B22R-like proteins, ring finger host-range proteins, virion-associated proteins, glycoproteins and transcription-related proteins, among others. Regarding coding-region differences between Italian and Spanish strains, two in-frame deletions were shared by the Spanish genomes but absent from the Italian genomes: a polymerase subunit and a kelch-like protein. Complete variant annotations are provided in **supplementary Table 2**.

**Figure 3.**
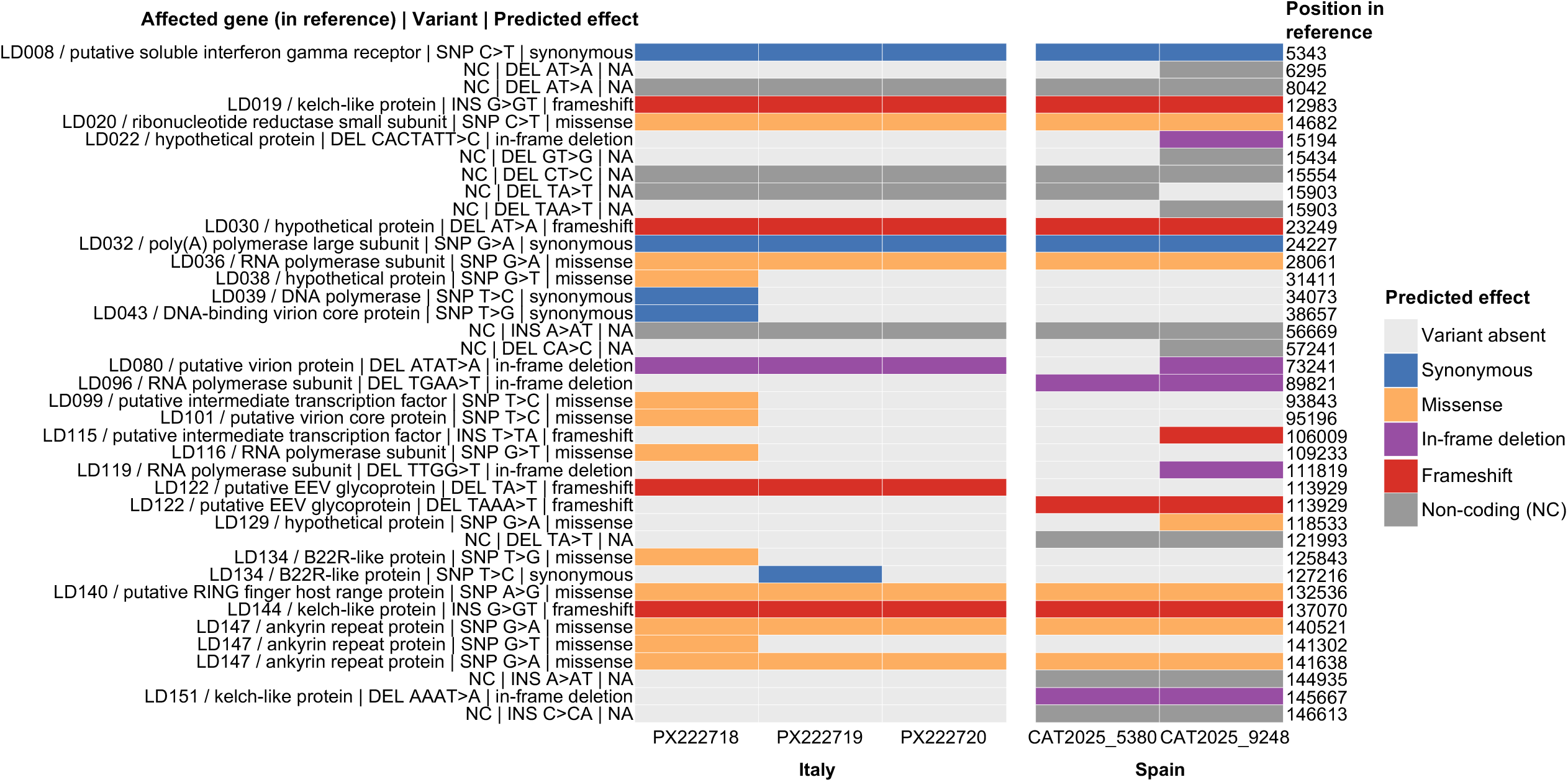
Variations detected in south-western European available strains (Spain and Italy) using the Nigerian V281 strain as reference. The reference genome position, the affected gene, the type of change and the predicted effect are detailed. The colour code corresponds to the predicted effect of each variation. NC (non-coding) indicates that the variant does not affect a coding region.

## Discussion

In this study, we report the genomic characterization of two LSDV isolates associated with the first confirmed outbreak in Catalonia, north-eastern Spain. The two genomes showed a high degree of similarity, supporting their association with the same outbreak strain. The genomic analysis provides relevant information to assess the relationship between these strains and recently reported European and African strains. Previously, Italy confirmed its first outbreak of LSDV in Orani (Sardinia) on 21 June 2025, whereas France reported its first case in Chambéry (Savoie) on 29 June 2025.

The phylogenetic reconstruction confirmed the major LSDV genomic structure previously observed (30,31), clearly separating subgroup 1.1, corresponding to the Neethling-like lineage, from subgroup 1.2, which includes Kenya-like and recent wild-type viruses. Within subgroup 1.1, our analysis separated South-African from Asian (R4 lineage) and Russian strains, as described in the literature. Within subgroup 1.2, the topology reproduced the main subdivision between clade 1.2a, comprising East African and Asian-related viruses, and clade 1.2b, which includes recent wild-type viruses from southern Africa, the Middle East, south-eastern Europe and Central Asia. The two genomes analysed in this study were placed within clade 1.2 and clustered with the Sardinian 2025 strains and the Nigerian isolate V281, rather than with strains from previous outbreaks in south-eastern Europe. Thus, the present analysis supports the inclusion of the Catalonian outbreak strains within the Italy/Nigeria-associated group of the clade 1.2, consistent with the topology recently reported for the Italian outbreak (15). This placement was also consistent with the *rpo30*-based phylogeny, which allowed a broader representation of African diversity than the whole-genome dataset. Despite the limited resolution of this single-gene marker, and the non-conclusive placement of the Cameroonian sequences, the analysis significantly clustered the Spanish and Italian sequences together with several Nigerian sequences, supporting their separation from the south-eastern European strains.

Additional complete genomes from central and north African strains would be essential to better investigate the relationship between the recent European outbreaks and African LSDV diversity. In fact, recent gene-based analyses of the first Algerian LSDV outbreak in 2024 also suggest a possible North African connection, as Algerian strains appeared close to recent Italian strains in some phylogenetic inferences (32). Although these sequences were not available at the time of our analysis, they reinforce the need for additional complete genomes from North Africa to assess whether this region may have contributed to the recent introduction of LSDV into south-western Europe. Similarly, two recent reports have also documented LSDV cases in North Africa (38,39), although neither provided publicly available sequencing data associated with these outbreaks. One of them included a *p32* gene-based phylogeny, in which the Algerian and Tunisian amplicons clustered with other African and Asian LSDV sequences (38). Considering that the analysis was based on a restricted country dataset and the authors’ discussion that this marker is “limited and poorly discriminative”, these data do not allow a robust assessment of the phylogenetic placement of these strains. Nevertheless, the observed clustering pattern may still suggest the possible circulation of different LSDV lineages in North Africa, including viruses related to Asian strains.

The comparison of variants using the 2018 Nigerian V281 strain as reference was performed because this strain represented the closest available complete African genome within the same clade. Therefore, it provided a useful reference to explore which mutational changes may have been accumulated in the recent south-western European LSDV genomes. Overall, the Spanish and Italian genomes showed a highly similar variant profile, with many changes shared between both countries, suggesting a conserved pattern. Interestingly, several non-synonymous changes affected genes in which variability is commonly observed in LSDV and other capripoxviruses, and which have often been discussed in relation to host interaction, host range, immune modulation or virulence-related traits. These included genes encoding kelch-like, ankyrin-repeat, B22R-like and ring finger host-range proteins, as well as some virion-associated, glycoprotein and transcription-related proteins (30,40,41). Despite the overall similarity between countries, the Spanish genomes differed from the Italian genomes by two shared coding changes affecting a kelch-like protein and an RNA polymerase subunit, both of which could be considered of interest given their potential involvement in host interaction, viral replication or virulence-related phenotypes, as previously described (30,40). These country-associated variants could suggest that the virus has continued accumulating changes during or between the recent European outbreaks. Nevertheless, this interpretation should remain cautious, particularly for variants detected in a single genome, as some differences may also be influenced by sequencing quality, assembly uncertainty or local alignment artefacts. In any case, the actual functional impact of the variants found in south-western Europe strains cannot be determined from sequence data alone, needing further investigation.

In conclusion, the genomic data presented here indicate that the LSDV strains detected in Catalonia belong to clade 1.2 and are closely related to the Mediterranean strains recently reported in Sardinia, Italy. These findings support a phylogenetic connection between the Spanish outbreak and the Italy/Nigeria-associated group, possibly through northern Africa, and highlight the need for additional complete genomes, to better resolve the origin and connection between recent LSDV outbreaks. Additional complete genomes from recent European and African outbreaks, particularly from France and North Africa, will be essential to clarify the origin and transmission pathways of LSDV incursions in the Mediterranean region.

## Supplementary materials

**Supplementary Figure 1. Maximum likelihood phylogenetic tree of all available *rpo*30 genes from LSDV**. The colour code follows the continent or geographical area of isolation, with Nigeria/Cameroon and Italy/Spain differentiated within each continent. The tree is rooted at midpoint. Yellow circles indicate branch bootstrap support equal or greater than 70. Branches with bootstrap support□<□70 have been collapsed. The clade of interest is highlighted in green.

**Supplementary Table 1. A)** Complete LSDV genomes included in the WGS-phylogenetic analysis. **B)** *rpo*30 genes retrieved from NCBI for gene-based phylogenetic reconstruction.

**Supplementary Table 2.** List of all variants found in each of the Spanish and Italian genomes using the Nigerian v281 strain as reference.

## Notes

### Competing Interest Statement

The authors have declared no competing interest.

