## Supplementary figures and images for "First outbreak of Lumpy Skin disease in Catalonia, Spain, 2025-2026"

### Supplementary figure 1

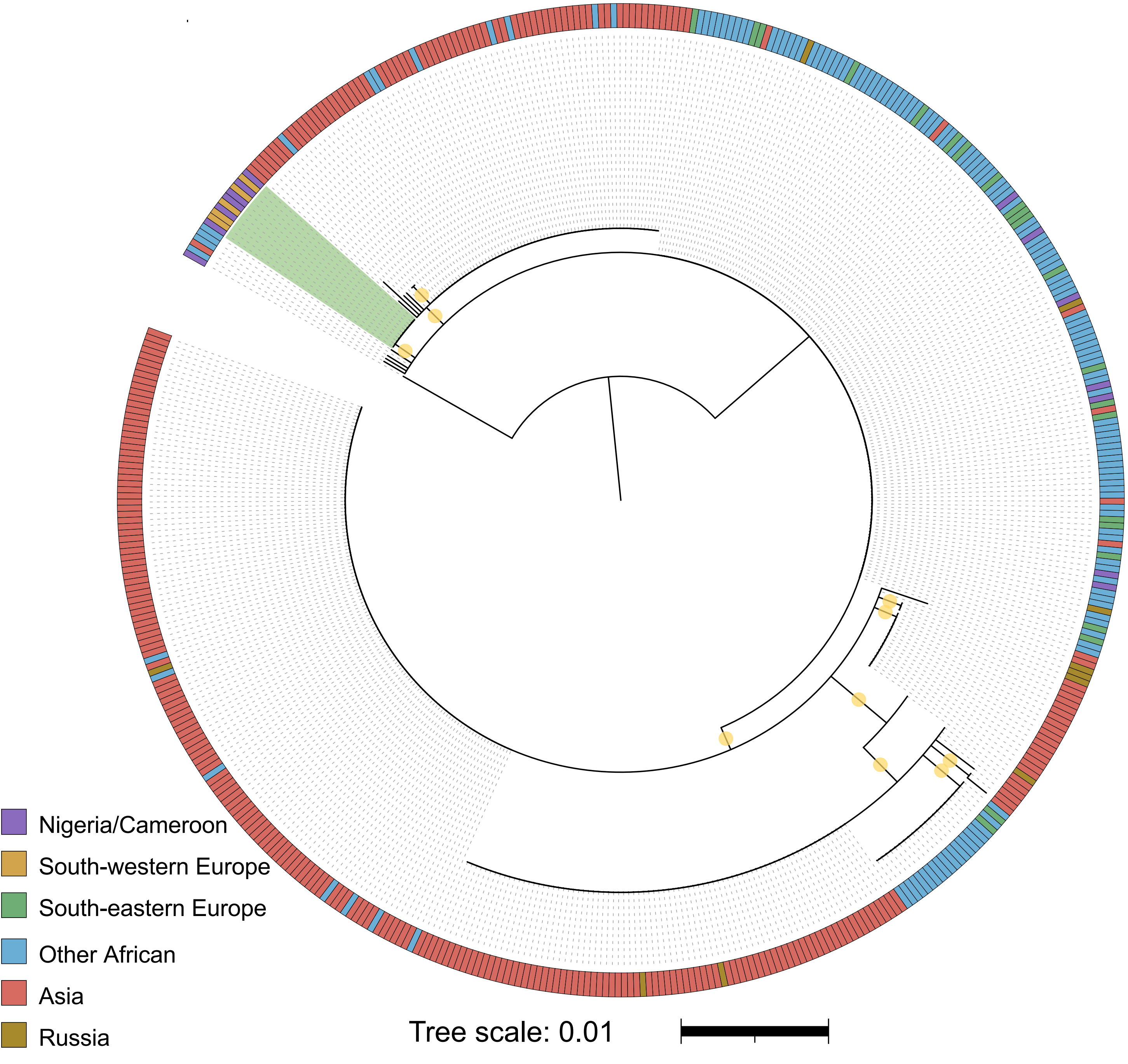
